# Inhibition of SARS-CoV-2 3CLpro *in vitro* by chemically modified tyrosinase from Agaricus bisporus

**DOI:** 10.1101/2023.03.13.532357

**Authors:** David Aguilera, David Ortega-Alarcon, Olga Abian, Adrian Velazquez-Campoy, Jose M. Palomo

**Affiliations:** Instituto de Catálisis y Petroleoquímica (ICP), CSIC, C/ Marie Curie 2. 28049 Madrid; Institute of Biocomputation and Physics of Complex Systems (BIFI), Joint Unit GBsC-CSIC-BIFI, University of Zaragoza, Zaragoza, 50018, Spain; Fundación Instituto de Investigación Sanitaria de Aragón (IIS Aragón), 50009 Zaragoza, Spain; CIBERehd, 28029 Madrid, Spain

**Keywords:** *Agaricus bisporus*, tyrosinase, SARS-CoV-2, COVID-19, 3CLpro, therapeutic proteins, protein modification, enzymes

## Abstract

Antiviral compounds are crucial to controlling the SARS-CoV-2 pandemic. Approved drugs have been tested for their efficacy against COVID-19, and new pharmaceuticals are being developed as a complementary tool to vaccines However, there are not any effective treatment against this disease yet. In this work, a cheap and fast purification method of natural tyrosinase from *Agaricus bisporus* fresh mushrooms was developed in order to evaluate the potential of this enzyme as a therapeutic protein by the inhibition of SARS-CoV-2 3CLpro protease activity *in vitro*. Tyrosinase showed a mild inhibition of 3CLpro of around 15%. Thus, different variants of this protein were synthesized through chemical modifications, covalently binding different tailor-made glycans and peptides to the amino terminal groups of the protein. These new tyrosinase conjugates were purified and characterized by circular dichroism and fluorescence spectroscopy analyses, and their stability under different conditions. Then all these tyrosinase conjugates were tested in 3CLpro protease inhibition. From them, the conjugate between tyrosinase and dextran-aspartic acid (6kDa) polymer showed the highest inhibition, with an IC_50_ of 2.5 μg/ml and IC_90_ of 5 μg/ml, results that highlight the potential use of modified tyrosinase as a therapeutic protein and opens the possibility of developing this and other enzymes as pharmaceutical drugs against diseases.

## 1. Introduction

The novel coronavirus SARS-CoV-2 emerged in the end of 2019 in Wuhan, China, causing a disease that has spread worldwide named COVID-19, declared as a pandemic by the World Health Organization (WHO) in early 2020 (Hu et al., 2021). To date, no antivirals or therapy has been proven effective to treat this disease (Jamalipour Soufi and Iravani, 2020). Consequently, many research efforts are put into finding treatments, and developing vaccines, which has been successful with several already in the market (Forni et al., 2021). Potential therapeutic drugs are aimed to target proteins involved in essential processes for the viral infection and replication. This includes proteins from the cellular host, including the receptor Angiotensin-converting enzyme-related carboxypeptidase (ACE2) or the transmembrane protease serine 2 (TMPRSS2), and structural viral proteins such as the spike protein (Hu et al., 2021)Non-structural viral proteins are involved in the genome and structural protein replication and thus constitute a target for inhibitory drugs (Zhang and Tang, 2021). Among these, the RNA dependent RNA polymerase (RdRp) and the 3-chymotrypsin like protease (3CLpro) have been a focus for the development of new drugs, because they are essential for viral replication and there are no similar targets in the host organism (Anand et al., 2002).SARS-CoV-2 contains a large positive single-stranded RNA genome, that codifies for the structural proteins and two overlapping polyproteins (pp1a and pp1ab). 3CLpro is responsible for the processing of the C-terminal region of these polyproteins, including 11 non-structural proteins (Hegyi and Ziebuhr, 2002; Jin et al., 2020). There are two types of 3CLpro potential inhibitors: repurposing of already approved drugs, and the use of small molecules or peptides as potential inhibitors, which can be found by high-throughput methods (Zhang and Tang, 2021).

Tyrosinase (EC 1.14.18.1) is an enzyme found in an extensive number of species, including human, animals, plants, bacteria and fungi. In the case of *Agaricus bisporus*, six different genes codify for tyrosinase isoforms (*ppo1-6*), of which *ppo3* and *ppo4* have been isolated from fruiting bodies (Ismaya et al., 2011; Pretzler et al., 2017). This enzyme contains two copper atoms in its active site, catalysing the hydroxylation of monophenols and the subsequent oxidation of the diphenols into quinones (Figure 1B) (Wichers et al., 2003). Tyrosinase is usually found in a 120 kDa tetrameric form, consisting of two subunits H of ~ 45 kDa each, and two subunits L of ~ 14 kDa. Only H subunits have been reported to have catalytic activity, and are known to be sufficient for this activity (Figure 1A) (Wichers et al., 1996).

**Figure 1.**
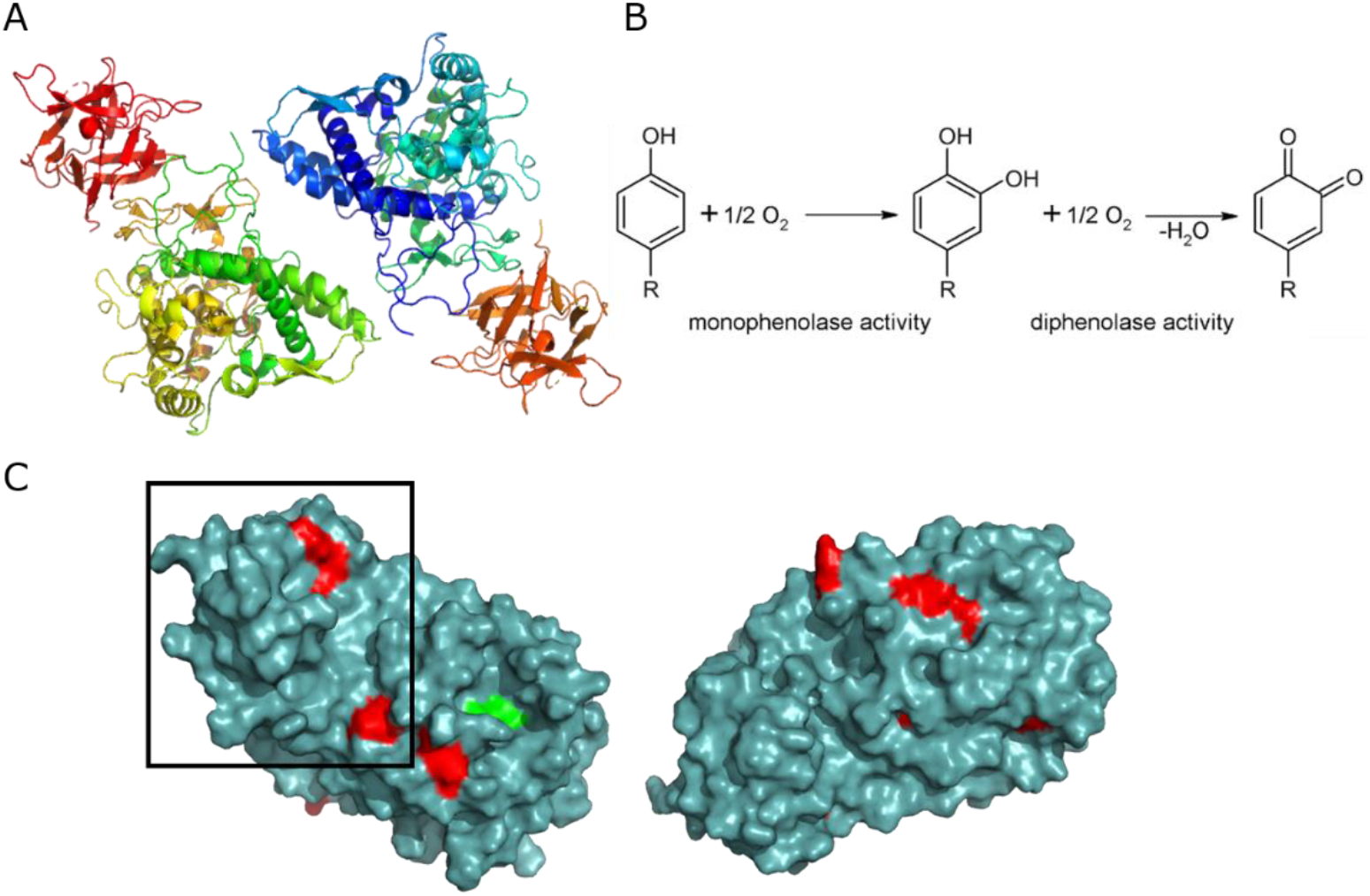
**A.** Structure of the tetrameric form of AbTyr. Protein Data Bank code: 2Y9W, image created using Pymol. **B.** Scheme of reactions catalysed by tyrosinase (Panis et al., 2020, CC by 4.0). **C.** Structure of 3CLpro. Protein Data Bank code: 7ALH, image created using Pymol. Red: tyrosine residues. Green: active site. The zone in which the protein binds to form the homodimer is marked with a black square.

Recently, tyrosinase from *A. bisporus* (AbTyr) has been shown to have antiviral activity against the Hepatitis C virus (HCV). Enzymatic activity of AbTyr was proved to be responsible for the antiviral activity, as inactivated protein did not inhibit viral replication (Lopez-Tejedor *et al*., 2020). This inhibition may be caused by the hydroxylation of critical tyrosine residues of the HCV protease.

Herein, we hypothesize that AbTyr may also show inhibition of 3CLpro from SARS-CoV-2 by the hydroxylation of tyrosine residues nearby the active site of the viral protease or by disrupting the formation of the homodimer, crucial for its enzymatic activity (Figure 1) (Kneller et al., 2020). Post-translational protein modifications (PTMs) include those that naturally occur in the cells, such as ubiquitination, phosphorylation, methylation and acetylation, and also the introduction of different moieties, peptides or polymers by chemical modifications, usually with the aim to improve properties of the protein or add functions like fluorophores or tags (Boutureira and Bernardes, 2015; Stephanopoulos and Francis, 2011). PTMs can also be applied in the development of therapeutic proteins, for example the use of polyethylene glycol to reduce immunogenicity and increase the time the therapeutic protein circulates in the bloodstream (Cho et al., 2011).

In this work, these modifications were used to potentially enhance the inhibition of 3CLpro by AbTyr, by chemically binding glycans and peptides to terminal amine groups of this enzyme.

## 2. Materials and methods

### 2.1. Tyrosinase extraction from Agaricus bisporus

Mushrooms were purchased from supermarkets Mercadona (Champinter), Hiber (Neofungi) and Simply (Laumont). The protocol for the tyrosinase extraction was the following: 15 g of mushrooms without the stipes were chopped and put under agitation in 50 ml of cold acetone for 30 min in the Bunsen rod stirrer AGV-8 and centrifuged at 7000 rpm and 4°C for 20 min using the centrifuge Biocen 22R. The supernatant was discarded, and the mushrooms were put under agitation in 50 ml of cold water for 1 h in the Nahita DJ-1 rod stirrer. The mix was centrifuged at 8000 rpm 4°C 20 minutes, the pellet was discarded and 15g of ammonium sulfate (Sigma-Aldrich) were slowly added to the supernatant under gentle agitation using a magnet stirrer. This protein extract was centrifuged at 8000 rpm 4°C for 40 min, and the pellet was stored at −20°C. Dialysis was performed using a tubing cellulose membrane (D9652, molecular weight cut-off = 14 kDa, Sigma-Aldrich) in 1L water or sodium phosphate 25 mM buffer at pH 7 for 30 min, repeated 3 times. Protein solutions after dialysis were stored at 4°C.

### 2.2. Tyrosinase enzymatic activity assay

The enzymatic activity of tyrosinase was measured in the presence of 1 ml of L-3,4-dihydroxy-phenylalanine (L-DOPA, Sigma-Aldrich) 1 mM in sodium phosphate buffer 0.1 M pH 7 at room temperature. The increase in absorbance at 475 nm after adding 25 μL of sample was measured using the V-730 spectrophotometer (Jasco). An increase of 0.001 of absorbance in 1 minute was defined as an enzymatic activity unit (U). Enzymatic activity was also tested in the presence of L-ascorbic acid or hydroquinone 1 mM (Sigma-Aldrich), or acetonitrile 40% (v/v).

### 2.3. SDS-PAGE electrophoresis

Sodium dodecyl sulphate–polyacrylamide gel electrophoresis (SDS-PAGE) was performed using the system PerfectBlue™ Double Gel System Twin S from Peqlab, and the PS300B Volt Power Supply from Hoefer at 170 V and constant amperage. 12% resolving gel (acrylamide 12%, 0.375 M Tris-HCl pH 8.8, 0.1% SDS, 0.05% APS, TEMED) and 5% stacking gel (acrylamide 5%, 0.136 M Tris-HCl pH 6.8, 0.1% SDS, 0.05% APS, TEMED) were used and was run in a buffer 0.2 M glycine, 0.25M Tris-HCl pH 8.8, 1% SDS. The marker used for the gels was Unstained Protein Molecular Weight Marker from Thermoscientific. Gels were stained with Coomassie Brilliant Blue 0.25% and unstained with a solution of 43% methanol and 7.5% acetic acid. Samples were prepared in a 1:1 solution with 2x loading buffer (80 mM Tris-HCl pH 6.8, 4% SDS, 20% glycerol, 10% 2-mercaptoethanol, Bromophenol Blue 0.02%). Silver staining was performed following the Protocol for Silver Staining of Gels from Alphalyse, using sodium sulfate instead of sodium thiosulfate.

### 2.4. Protein adsorption on solid supports

Tyrosinase adsorption was tested on Butyl-Sepharose (C4), Octyl-Sepharose (C8, from GE Healthcare), geranyl-functionalized carboxymethylcellulose (G-CM), Octadecyl Sepabeads (C18, from RESINDION) and Epoxy-C18 (E-C18, from Purolite). The adsorption was performed using 4 ml of sample 0.4 mg / ml or 0.25 mg / ml with 0.3 g of solid support, in water or in sodium phosphate 25 mM or 100 mM pH 7 in a roller for 1 h. Different concentrations of Triton X-100 (0.01% - 0.25%) were used for the desorption of the proteins from the solid phase after an 1 h incubation.

### 2.5. Covalent binding of polymers and peptides

Proteins were modified using the following peptides Ac-DD, Ac-DEGD, Ac-FFD, Ac-AGAG, Ac-FDLG and Ac-AAGTA from GenScript, and the polymers containing carboxylic groups: commercial polymers: hyaluronic acid (8 – 15 kDa), polygalacturonic acid (25 – 30 kDa) and polyethyleneimine 25 kDa from Sigma-Aldrich, and tailor-made synthesized Dextran-Aspartic 6 kDa, Dextran-Aspartic 2 MDa, Dextran-Lysine 6 kDa and Dextran-Glycine 6 kDa (Romero et al., 2013). The different polymers were incubated in the presence of 10 eq N-ethyl-N’-(3- (dimethylaminopropyl)carbodiimide (EDC) (Tokyo Kasei) and 15 eq N-hydroxysuccinimide (NHS) (Sigma-Aldrich) in water at pH 4.5 for 1 h. 50 or 100 eq of activated polymers were incubated with 1.6 mg of proteins supported on a solid phase of C18 (0.3 g) overnight in sodium phosphate 50 mM or water at pH 7 and then washed with sodium phosphate buffer 25 mM pH 7. Modified proteins were eluted from the support using 0.05 % or 0.1 % Triton X-100.

### 2.6. 3CLpro inhibition assay

*In vitro* 3CLpro catalytic activity was determined using a fluorescence resonance energy transfer (FRET) assay with the peptide (Dabcyl)-KTSAVLQSGFRKME-(Edans)-NH2 (Biosyntan GmbH) as substrate. A concentration of 0.2 μM of protease was used in sodium phosphate 50 mM, NaCl 150mM pH 7, and the enzymatic reaction was started adding the peptide up to 20 μM, in a volume of 100 μL. The microplate reader FluoDia T70 (Photon Technology International) was used to continuously measure fluorescence for 60 minutes at an excitation wavelength of 380 nm and emission length of 500 nm. Protease activity was quantified as the increase in fluorescence. Inhibition of this activity was defined as the difference between protease activity in the presence or absence of the inhibitor and was evaluated by adding up to 400 μg / mL of these compounds.

### 2.7. Circular dichroism and fluorescence spectroscopy

Circular dichroism of the tyrosinase samples was measured using Chirascan spectropolarimeter (Applied Photophysics) at 25 °C. Far-UV spectra was recorded in a 0.1 cm path-length cuvette at wavelengths between 200-260 nm and a protein concentration of 10 μM in phosphate buffer saline at pH 7.2. Fluorescence spectroscopy measurements of 2 μM of each protein sample were obtained using the Varian Cary Eclipse Fluorescence Spectrophotometer (Agilent Technologies) with an excitation wavelength of 280 nm and recording the emission spectra between 300 nm and 400 nm.

## 3. Results and Discussion

### 3.1. Protein extraction from mushrooms

Mushrooms for the protein extraction following the protocol previously described were bought from four different sources, with the goal of analysing potential differences between batches and comparing their yield and enzymatic activity (Table 1). The presence of AbTyr and other proteins in the extracts was validated with an SDS-PAGE (Figure 2).

**Table 1:** Protein yield and specific diphenolase activity obtained from mushrooms.

| Mushrooms | Protein yield (mg) | Specific activity (U/μg) |
| --- | --- | --- |
| Champinter | 108 | 1.81 |
| Neofungi | 121 | 3.63 |
| Laumont (var. brunnescens) | 182 | 3.53 |
| Laumont | 160 | 3.23 |

**Figure 2.**
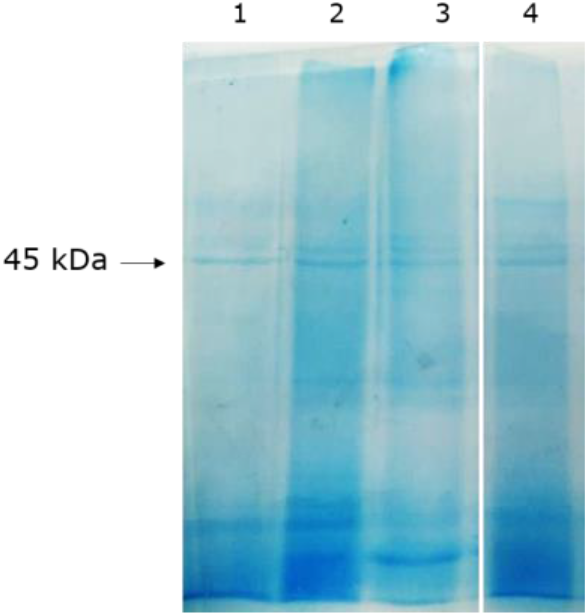
SDS-PAGE gel of mushrooms protein extracts. 1: Champinter, 2: Neofungi, 3: Laumont (var. brunnescens), 4: Laumont

AbTyr from Champinter presents a lower specific activity than the other extracts. This difference can be explained by a smaller band in the electrophoresis gel at 50 kDa, which corresponds to an AbTyr highly active isoform (Lopez-Tejedor and Palomo, 2018). Previously, inhibition of 3CLpro by AbTyr *in vitro* was tested and protease activity remained at around 85% of its activity without the tyrosinase. 3CLpro inhibition activity by these four extracts were tested. However, this experiment showed that the samples exhibited intrinsic protease activity, distorting the fluorescence of the FRET assay and highlighting the necessity of further purifying the tyrosinase in order to remove proteases from the mushroom extracts. Mushrooms from Neofungi were selected to perform the following experiments, as their protein extract presented a higher specific activity of abTyr.

### 3.2. Development of a tyrosinase purification method

Purification of AbTyr from the mushroom extract was attempted by binding it to several hydrophobic supports, composed of carbon chains of different length from 4 to 18 carbon groups. The adsorption was tested using the extract directly or by first dialyzing the proteins in sodium phosphate buffer 25 mM with the purpose of removing ammonium sulfate present from the protein precipitation step of the extraction protocol. While in most supports the absorption did not occur in a percentage higher than 40%, in the case of C18 sodium phosphate buffer 0.1 M pH 7, an 80% of offered protein was bound to the solid phase. Interestingly, when using C8, almost all enzymatic activity remained in the supernatant, while other proteins were retained in the support, as shown in the SDS-PAGE (Figure 3A).

**Figure 3.**
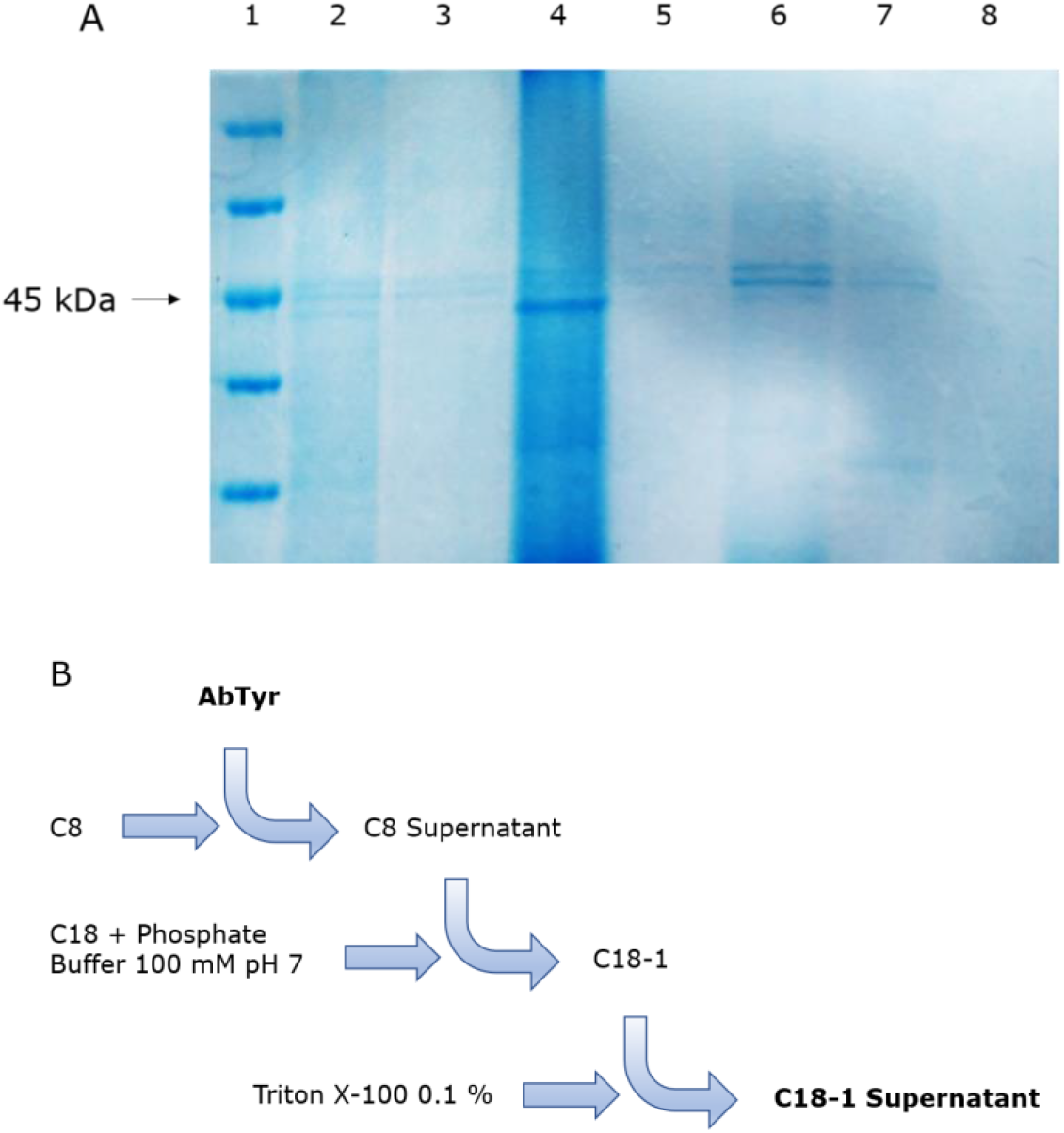
**A.** SDS-PAGE of AbTyr purification cascade. 1: Marker. 2: Extract. 3: C8 Supernatant. 4: C8. 5: C18 supernatant. 6: C18. 7: Triton X-100 elution supernatant. 8: Triton X-100 C18. **B.** Scheme of AbTyr purification cascade. C18-1: Proteins supported on C18.

A purification method was finally developed to obtain AbTyr purified and bound to a solid phase or eluted. It consists of an incubation with C8 of 4 mL 0.4 mg / mL of dialyzed protein in sodium phosphate buffer 25 mM pH 7 for 1 hour, and a consequent incubation of the supernatant with C18 in sodium phosphate buffer 100 mM pH 7 for 1 hour. Bound AbTyr can be eluted using 0.1% Triton X-100 (Figure 3B).

After the purification cascade, the final AbTyr sample contains around 50% of its initial enzymatic activity, and if incubated in C18 for 24 hours, it would retain around 30% of the initial activity.

### 3.3. Solid-phase protein modifications using the EDC/NHS strategy

With the objective of enhancing the inhibitory activity of AbTyr against 3CLpro, several modifications were performed using different glycans and peptides. These polymers may help AbTyr to recognize 3CLpro tyrosine residues by allosteric reasons and / or because of charges interacting with nearby amino acids.

The covalent linking is targeted at the free amine groups of the tyrosinase. This is performed using the EDC/NHS strategy, by which the carboxylic groups of the polymers are activated by EDC, and then stabilized by NHS, allowing them to react with the amine groups of the enzyme forming a covalent bond. As glycans, tailor-made dextran aldehyde (2000 kDa or 6 kDa) modified with aspartic acid, lysine or glycine (Romero et al., 2013), hyaluronic acid, and polygalacturonic acid were used. In the case of peptides, several containing mainly negatively charged (aspartic acid and glutamic acid residues) and / or hydrophobic amino acids (alanine, glycine and phenylalanine) were used.AbTyr before the last step of elution (Figure 3) was used to perform the post-translational modifications, as the solid phase makes this process faster and easier, avoiding extra steps to separate the enzyme from free excess polymers. All tyrosinase conjugates were then tested for their enzymatic activity. Most of them retained a similar diphenolase activity, except for Tyr-Polygal which lost its activity. Their stability in different temperatures or the presence of inhibitors or organic solvents were also evaluated. Significant changes were found in the case of Tyr-DextAsp-2000kDa, which enzymatic activity decreased importantly at 45 °C compared to AbTyr. When exposed to acetonitrile 40% (v/v), AbTyr lost its activity, while some modifications retained activity: Tyr-DextAsp-6kDa and Tyr-DGED a 5%; Tyr-Hyal, Tyr-FDLG and Tyr-AAGTA a 15-20%, and Tyr-DD a 40% of its initial enzymatic activity. Circular dichroism and fluorescence spectroscopy of the samples was measured to compare the shifts with unmodified AbTyr samples (Figure 4).

**Figure 4.**
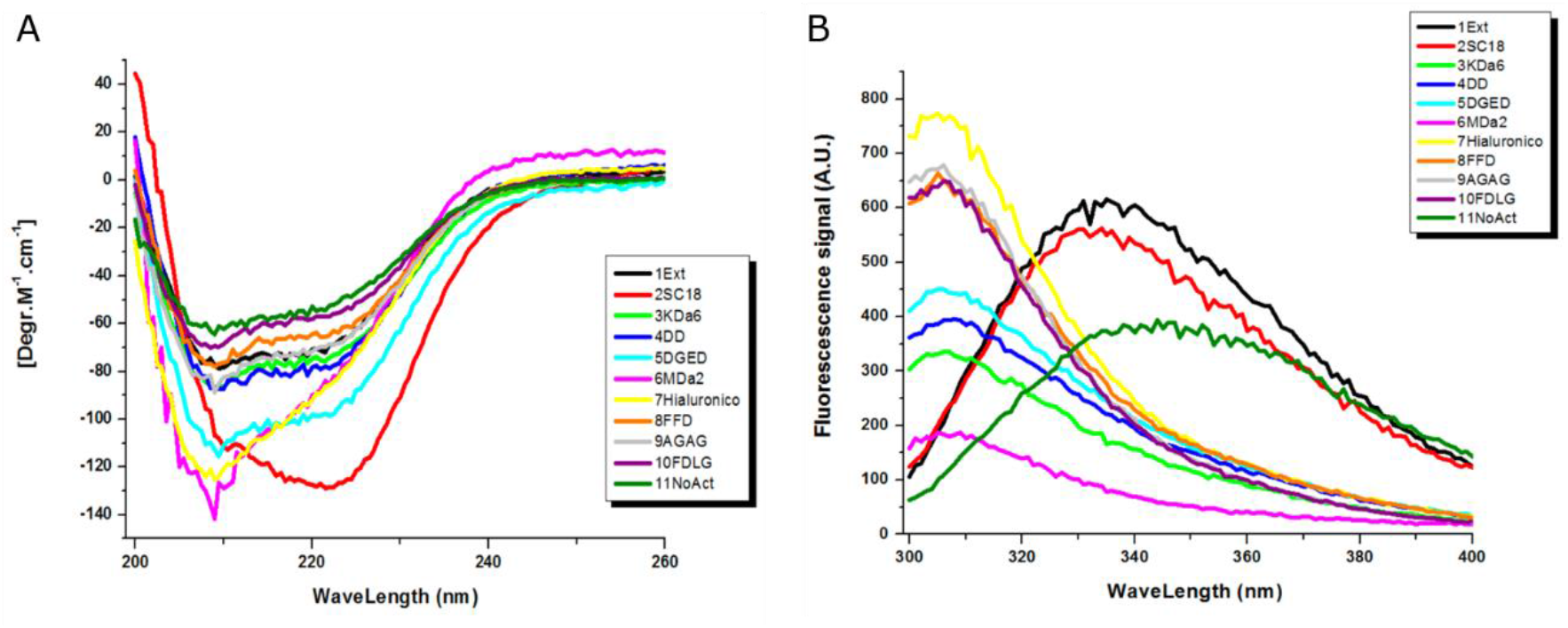
**A.** Circular dichroism of AbTyr samples. **B.** Fluorescence spectroscopy of AbTyr samples. SC18: C18 supernatant. NoAct: AbTyr without enzymatic activity after heat deactivation.

### 3.4. 3CLpro inhibition by AbTyr

AbTyr after purification and with its modifications was tested as an inhibitor of 3CLpro protease activity. Remaining intrinsic protease activity of the samples had to be taken into consideration when obtaining the inhibitory concentrations, by subtracting this activity to the present when added to 3CLpro. In Table 2 are represented the inhibitory concentrations of sample at which 3CLpro protease activity is reduced to a 50% (IC_50_) and to a 90% (IC_90_).

**Table 2.** Inhibitory concentrations of AbTyr against 3CLpro.

| Polymer | IC<sub>50</sub> (µg / ml) | IC<sub>90</sub> (µg / ml) |
| --- | --- | --- |
| Tyr-DextAsp-6kDa | 2.5 | 5 |
| Tyr-DextAsp-2MDa | - | - |
| Tyr-DextLys-6kDa | - | - |
| Tyr-DextGly-6kDa | - | - |
| Tyr-Hyal | 10 | 40 |
| Tyr-Polygal | 5 | 20 |
| Tyr-DD | 17.5 | >40 |
| Tyr-DGED | 35 | >40 |
| Tyr-FFD | 7.5 | 15 |
| Tyr-AGAG | 40 | >40 |
| Tyr-FDLG | - | - |
| Tyr-AAGTA | - | - |

Tyrosinase modifications with cationic polymers did not present significative inhibition of 3CLpro activity, while carboxylated polymers appear to present better results. The best result obtained was by Tyr-DextAsp-6KDa, with an IC_50_ of 2.5 μg / ml and IC_90_ of 5 μg / ml of sample (Figure 5). Other conjugates that also present significant inhibition *in vitro* were Tyr-Hyal with an IC_50_ of 10 μg / ml and IC_90_ of 40 μg / ml, and Tyr-Polygal with an IC_50_ of 5 μg / ml and IC_90_ of 20 μg / ml. This last one was surprising, as this modification cause the loss of diphenolase activity: the positive result as a 3Clpro inhibitor might be explained by its monophenolase activity. Other promising results were showed by peptides DD and FFD (Table 2).

**Figure 5:**
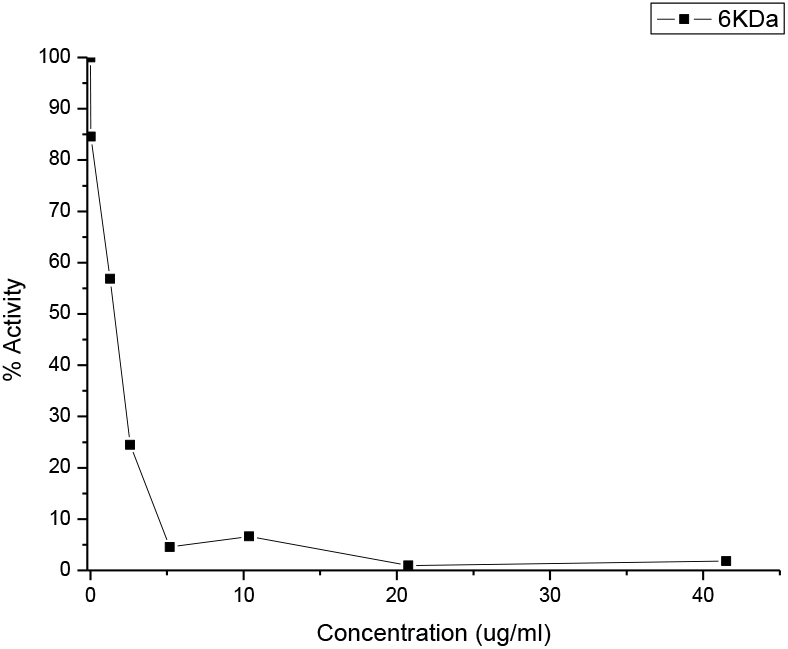
Inhibitory activity of Tyr-DextAsp-6kDa against 3CLpro protease activity.

Free polymers were also tested for inhibition of 3CLpro activity, obtaining negative results confirming that this inhibition was caused by the presence of the protein sample.

## 4. Conclusions

In this work, we developed a new and fast purification method of *Agaricus bisporus* tyrosinase after its extraction from mushrooms. This enzyme showed a mild inhibitory activity of 3CLpro protease activity *in vitro*. We also developed new tyrosinase bioconjugates with different glycans and peptides using a solid phase which were proved to enhance the inhibitory activity of tyrosinase against 3CLpro, obtaining an IC_50_ of 2.5 μg / ml and IC_90_ of 5 μg / ml of sample in the case of the use of the glycan Dextran 6 kDa with aspartic acid groups (Tyr-DextAsp-6kDa).

These results highlight the potential capability of enzymes as therapeutic proteins, in particular of tyrosinase against SARS-CoV-2 3CLpro, adding to the recent research on the activity of this protein against HCV. Further steps need to be performed in order to assess the potential of modified AbTyr as a pharmaceutical drug, including cytotoxicity tests, *in vivo* assays and evaluating its ability to enter the cellular membrane.

## Acknowledgements

This work was supported by the Spanish National Research Council (CSIC) (project CSIC PTI-Global Health SGL2103036 (J.M.P). This work was supported by Fundación hna (grant to A.V.-C. and O.A.).

## References

1. Anand, K., Palm, G.J., Mesters, J.R., Siddell, S.G., Ziebuhr, J., and Hilgenfeld, R. (2002). Structure of coronavirus main proteinase reveals combination of a chymotrypsin fold with an extra α-helical domain. EMBO J. 21, 3213–3224.

2. Boutureira, O., and Bernardes, G.J.L. (2015). Advances in chemical protein modification. Chem. Rev. 115, 2174–2195.

3. Cho, H., Daniel, T., Buechler, Y.J., Litzinger, D.C., Maio, Z., Putnam, A.M.H., Kraynov, V.S., Sim, B.C., Bussell, S., Javahishvili, T., et al. (2011). Optimized clinical performance of growth hormone with an expanded genetic code. Proc. Natl. Acad. Sci. U. S. A. 108, 9060–9065.

4. Forni, G., Mantovani, A., Forni, G., Mantovani, A., Moretta, L., Rappuoli, R., Rezza, G., Bagnasco, A., Barsacchi, G., Bussolati, G., et al. (2021). COVID-19 vaccines: where we stand and challenges ahead. Cell Death Differ. 28, 626–639.

5. Hegyi, A., and Ziebuhr, J. (2002). Conservation of substrate specificities among coronavirus main proteases. J. Gen. Virol. 83, 595–599.

6. Hu, B., Guo, H., Zhou, P., and Shi, Z.L. (2021). Characteristics of SARS-CoV-2 and COVID-19. Nat. Rev. Microbiol. 19, 141–154.

7. Ismaya, W.T., Rozeboom, H.J., Weijn, A., Mes, J.J., Fusetti, F., Wichers, H.J., and Dijkstra, B.W. (2011). Crystal structure of agaricus bisporus mushroom tyrosinase: Identity of the tetramer subunits and interaction with tropolone. Biochemistry 50, 5477–5486.

8. Jamalipour Soufi, G., and Iravani, S. (2020). Potential inhibitors of SARS-CoV-2: recent advances. J. Drug Target 29(4), 349–364.

9. Jin, Z., Du, X., Xu, Y., Deng, Y., Liu, M., Zhao, Y., Zhang, B., Li, X., Zhang, L., Peng, C., et al. (2020). Structure of Mpro from SARS-CoV-2 and discovery of its inhibitors. Nature 582, 289–293.

10. Kneller, D.W., Phillips, G., O’Neill, H.M., Jedrzejczak, R., Stols, L., Langan, P., Joachimiak, A., Coates, L., and Kovalevsky, A. (2020). Structural plasticity of SARS-CoV-2 3CL Mpro active site cavity revealed by room temperature X-ray crystallography. Nat. Commun. 11, 1–6.

11. Lopez-Tejedor, D., and Palomo, J.M. (2018). Efficient purification of a highly active H-subunit of tyrosinase from Agaricus bisporus. Protein Expr. Purif. 145, 64–70.

12. Lopez-Tejedor, D., Clavería-Gimeno, R., Velazquez-Campoy, A., Abian, O., and Palomo, J.M. (2020). Tyrosinase from mushroom Agaricus bisporus as an inhibitor of the Hepatitis C virus. BioRxiv 2020.12.23.424187.

13. Panis, F., Kampatsikas, I., Bijelic, A., and Rompel, A. (2020). Conversion of walnut tyrosinase into a catechol oxidase by site directed mutagenesis. Sci. Rep. 10, 1–14.

14. Pretzler, M., Bijelic, A., and Rompel, A. (2017). Heterologous expression and characterization of functional mushroom tyrosinase (AbPPO4). Sci. Rep. 7, 1–10.

15. Romero, O., Rivero, C.W., Guisan, J.M., and Palomo, J.M. (2013). Novel enzyme-polymer conjugates for biotechnological applications. PeerJ 2013, e27.

16. Stephanopoulos, N., and Francis, M.B. (2011). Choosing an effective protein bioconjugation strategy. Nat. Chem. Biol. 7, 876–884.

17. Wichers, H.J., Gerritsen, Y.A.M., and Chapelon, C.G.J. (1996). Tyrosinase isoforms from the fruitbodies of Agaricus bisporus. Phytochemistry 43, 333–337.

18. Wichers, H.J., Recourt, K., Hendriks, M., Ebbelaar, C.E.M., Biancone, G., Hoeberichts, F.A., Mooibroek, H., and Soler-Rivas, C. (2003). Cloning, expression and characterisation of two tyrosinase cDNAs from Agaricus bisporus. Appl. Microbiol. Biotechnol. 61, 336–341.

19. Zhang, Y., and Tang, L. V. (2021). Overview of Targets and Potential Drugs of SARS-CoV-2 According to the Viral Replication. J. Proteome Res. 20, 49–59.

